# Mouse mammary tumor virus is implicated in severity of colitis and dysbiosis in the IL-10^-/-^ mouse model of inflammatory bowel disease

**DOI:** 10.1101/2023.01.26.525771

**Authors:** Heather Armstrong, Mandana Rahbari, Heekuk Park, David Sharon, Aducio Thiesen, Naomi Hotte, Ning Sun, Hussain Syed, Hiatem Abofayed, Weiwei Wang, Karen Madsen, Eytan Wine, Andrew Mason

## Abstract

**Background:** Following viral infection, genetically manipulated mice lacking immunoregulatory function may develop colitis and dysbiosis in a strain specific fashion that serves as a model for inflammatory bowel disease (IBD). We found that one such model of spontaneous colitis, the interleukin (IL)-10 knockout (IL-I0^-/-^) model derived from the SvEv mouse, had evidence of increased mouse mammary tumor virus (MMTV) viral RNA expression compared to the SvEv wildtype. MMTV is endemic in several mouse strains as an endogenously encoded betaretrovirus that is passaged as an exogenous agent in breast milk. As MMTV requires a viral superantigen to replicate in the gut associated lymphoid tissue prior to the development of systemic infection, we evaluated whether MMTV may contribute to the development of colitis in the IL-10^-/-^ model.

**Results:** Viral preparations extracted from IL-10^-/-^ weanling stomachs revealed augmented MMTV load compared to the SvEv wildtype. Illumina sequencing of the viral genome revealed that the two largest contigs shared 96.4% - 97.3% identity with the *mtv-1* endogenous loci and the MMTV(HeJ) exogenous virus from the C3H mouse. The MMTV *sag* gene cloned from IL-10^-/-^ spleen encoded the MTV-9 superantigen that preferentially activates T cell receptor Vβ-12 subsets, which were expanded in the IL-10^-/-^ versus the SvEv colon. Evidence of MMTV cellular immune responses to MMTV Gag peptides was observed in the IL-10^-/-^ splenocytes with amplified interferon-γ production versus the SvEv wildtype. To address the hypothesis that MMTV may contribute to colitis, we used HIV reverse transcriptase inhibitors, tenofovir and emtricitabine, and the HIV protease inhibitor, lopinavir boosted with ritonavir, for 12 weeks treatment versus placebo. The combination anti-retroviral therapy with known activity against MMTV was associated with reduced colonic MMTV RNA and improved histological score in IL10^-/-^ mice, as well as diminished secretion of pro-inflammatory cytokines and modulation of the microbiome associated with colitis.

**Conclusions:** This study suggests that immunogenetically manipulated mice with deletion of IL-10 may have reduced capacity to contain MMTV infection in a mouse-strain specific manner, and the antiviral inflammatory responses may contribute to the complexity of IBD with the development of colitis and dysbiosis.

## Introduction

Inflammatory bowel diseases (IBD) are debilitating chronic illnesses of the intestinal tract, thought to occur in genetically-predisposed individuals who are exposed to microbial, dietary and environmental triggers [1]. While eukaryotic viruses play a role in shaping mucosal immunity and homeostasis without causing pathology in healthy subjects [2], they can contribute to the development of IBD in genetically predisposed subjects by disturbing biophysical integrity of the bowel, altering the microbiome, provoking loss of tolerance to gut microbiota and promoting chronic inflammation [1, 3]. The microbial complexity of IBD is illustrated in metagenomic surveys of whole colon RNA samples, where increased expression of common viruses is associated with bacterial dysbiosis and an increased expression of endogenous retrovirus RNA, which effectively lowers the threshold for triggering inflammation through innate immune pathways [4]. However, studies of viral infection in IBD patients are limited by the difficulty in discerning a cause-and-effect relationship, differentiating passenger viruses from causal agents, and often do not account for the effects of specific IBD risk alleles.

Central to a viral hypothesis is that IBD is restricted to genetically susceptible individuals and involves alterations in the gut microbiome. Indeed, the genetic predisposition and pathogenesis of Crohn disease (CD) are intertwined with downregulation of innate immunity and autophagy pathways employed for handling of intracellular bacteria [5]. The superimposed role of viral infection in the IBD process is exemplified in “virus + susceptibility gene” animal models. For example, murine norovirus infection in the mutated *ATL16L1* mouse model triggers both enteritis and dysbiosis that is treatable by antibiotics [3]. The clinical relevance of this model is underscored by reports that norovirus infection mediates a critical loss of cytoprotective effect of Paneth cells leading to a CD-like pathology [6] and infection with norovirus may often precede the development of CD in population studies [7].

Both viral infection and candidate IBD susceptibility variants may also serve a conjoint effect of facilitating immunodeficiency that perturbs intestinal barrier function. Several retroviruses including HIV [8], simian immunodeficiency virus [9], feline leukemia virus [10], and murine leukemia virus [11] have all been linked with colitis with disruption of mucosal barrier function and dysregulation of immune responses to commensal microbes. Notably, human endogenous retroviral RNA is often overexpressed in inflammatory disease leading to activation of innate immune responses, but these agents do not emerge from the genome to make transmissible, exogenous viral particles [4]. In contrast, inbred laboratory mice have been selected to develop disease and as a result, genetically manipulated immunodeficient mice can resurrect endogenous retroviral sequences from the genome to create infectious transmissible particles that cause pathology [12].

We have studied this process and identified evidence of mouse mammary tumor virus (MMTV) infection in several genetically modified mice [13]. MMTV is the causal agent of murine breast cancer and has been linked with autoimmune biliary disease and murine rheumatoid arthritis in genetically manipulated models [13, 14]. MMTV is a murine betaretrovirus endogenously encoded in the genome of most laboratory mice. Some mice express transmissible betaretrovirus particles in a strain dependant fashion and develop breast cancer, whereas other mice do not develop productive infection due to modifications that inactivate replication competence of the endogenous *mtv* loci [14–16]. Furthermore, strains such as C57BL/6 have developed resistance to MMTV but infection may emerge following genetic manipulation of immune function [13]. Indeed, some of the MMTV expressing mice also serve as models for IBD, such as T cell TGF-β receptor II dominant-negative mouse and the IL-2 receptor α deficient mouse [13]. The natural biology of MMTV is germane to the pathophysiology of IBD. Neonatal infection in mouse milk occurs prior to the acidification of the stomach and virus is efficiently transmitted through mucosal surfaces by binding bacterial lipopolysaccharide and engagement of the toll-like receptor-4 (TLR-4) [14, 17]. This in turn leads to the production of interleukin (IL)-10 knockout (IL-10^-/-^) to tolerize the host to viral infection. A robust immune response is triggered in Peyer’s patches by the MMTV encoded viral superantigen (vSAG) to enable viral replication in proliferating lymphocytes [14]. The infected lymphocytes are trafficked to the mammary gland epithelium where MMTV is maximally expressed following pregnancy.

One commonly utilised research model of IBD is the IL-10^-/-^ mouse model of colitis, which not only resembles IBD but also responds to existing therapeutic options [18]. IL-10^-/-^ mice develop colitis shortly after weaning, which is highly dependent upon the environment the mice are raised in and the composition of their gut microbiome [19]. While evaluating different mouse models for evidence of MMTV [13], we incidentally found increased levels of MMTV RNA in tissues from the IL-10^-/-^ colitis model as compared to the 129 SvEv wildtype (WT) mice. The viral biology of MMTV may be relevant the IL-10^-/-^ colitis model because pups cross fostered with wild type mothers are protected from developing disease [20]. Furthermore, MMTV requires interaction with mucosal commensal bacteria to utilize the TLR-4 and IL-10 dependant mechanism for uptake and passage in gut-associated lymphoid tissue, while also dampening the antiviral immune responses [17].

The aim of this study was to address the hypothesis that MMTV infection contributes to development of spontaneous colitis observed in the IL-10^-/-^ model. Accordingly, we sought evidence of exogenous MMTV infection in weanling milk, vSAG activity in the colon, cellular immune responses to MMTV and studied the effects of combination antiretroviral therapy (cART) known to inhibit MMTV [21]. We observed a significant reduction in colonic histology scores in IL-10-/- mice, linked with reduced expression of MMTV RNA, pro-inflammatory cytokines within the colon and resultant changes in the microbiome in support of our hypothesis.

## Methods

### Animal Model

IL-10 deficient 129(B6)- IL-10^tm1Cgn^/J mice and wild-type mice on the same background were derived from DNAX Research Institute of Molecular and Cellular Biology Inc., Palo Alto, California and bred at the Health Sciences Laboratory Animal Services at the University of Alberta (Edmonton, Canada). All mice were kept under 25°C and 45-55% humidity conditions. The protocol was approved by the University of Alberta Health Research Ethics Board (Study ID. AUP0000510 and AUP0000138).

### Cellular immune response to MMTV Gag peptides

One million splenocytes from SvEv mice and IL-10^-/-^ mice (n=4) were seeded in duplicate in 24 well plates with 1 ml R10 medium (RPMI 1640 medium, 10% FBS, 1% Penicilin-Steptomycin, 25mM HEPES buffer and L-glutamine). Individual wells were stimulated for 24 hours at 37 °C with 2 μg/ml of 58 overlapping 15-20mer peptides covering the betaretrovirus Gag protein (Supplementary Table 1) synthesized by Mimotopes (Victoria, Australia); 50ng/ml PMA and 1 ug/ml Ionomycin served as a positive control, and the negative control “Nil” incorporated 30% DMSO used as a carrier for the peptides. After incubation for 24 hours at 37°C, 50 μl supernatant was assessed by ELISA to quantify interferon (IFN)-γ production using V-PLEX IFN-γ [range 0.04-570 pg/ml the Mesoscale Discovery (MSD) platform] following manufacturer’s directions.

### Next generation sequencing (NGS) and characterization of MMTV in IL10^-/-^ mouse milk

Mouse milk was obtained from stomachs of 3-5 day old IL-10^-/-^ and SvEv weanling pups (n=8), as described [22]. Lactoserum was derived by homogenizing stomachs in 2ml PBS, centrifuged at 2000g for 15 minutes at 4°C. Both the stomach pellet and lactoserum supernatant were used to make libraries. The lactoserum was further purified using a 30% PBS/sucrose cushion and centrifuged at 90,000g for 1 hour at 4°C. The lactoserum pellet was resuspended in 500μl of PBS and centrifuged at 10,000 RPM for 10 minutes at 4°C, the supernatant was processed through a 0.45μm filtered and stored at −80°C.

Total RNA was extracted from the lactoserum and stomach pellet using TRIzol reagent (Invitrogen), ribosomal RNA was removed using the Ribo-Zero rRNA removal kit (Illumina), converted to cDNA synthesis using superscript II (Invitrogen). MMTV nucleic acid was quantified using RT-qPCR, as described [13]. Total lactoserum cDNA was cloned into Illumina TruSeq libraries (RNAv2 kit, Illumina) and enriched for MMTV sequences using capture probes synthesised by Panomics for the Quantigene assay (RefSeq NC_001503.1). The capture was amplified by PCR for 10 cycles and sequenced using the MiSeq reagent kit v2 2×250 on an Illumina MiSeq instrument. For genome assembly, the sequences were assembled with the software SPAdes and then aligned against the RefSeq database of NCBI with Blast.

### Cloning of viral superantigen (vSAG)

The MMTV *sag* gene 3’ region was derived from IL-10^-/-^spleen RNA by conversion to complementary DNA using a SuperScript IV Reverse Transcriptase kit (Invitrogen), and PCR amplification with two separate primer sets: (i) SAg-F1: GGTGGCAACCAGGGACTTAT, Sag-R1: CCACTTGTCTCACATCCTCGT, Sag-F-inner1: CAACAGATGCCCCCTTACCA; and (ii) Sag-F5: AAAGAGGAGTGCGCTTGTCA, Sag-R5: ACCAAGTCAGGAAACCACTTGT, Sag-F-inner5: TTACAGACCAACAGACGCCC. Semi-nested PCR was conducted using 200ng of cDNA with Taq DNA Polymerase (Invitrogen) using thermocycler parameters set to 95°C: 5min [95°C: 35 seconds, 52°C: 30 seconds, 72°C 40 seconds] x 45 cycles. Both reactions provided two products per reaction that were gel purified in a 2% agarose gel using a GeneJet Gel Extraction kit (Thermo Scientific corp.). Each product was cloned using the pGEM®-T Easy vector kit (Promega Corp.) to derive 12 clones using the GeneJet plasmid purification kit (Thermo Scientific) for Sanger sequencing.

### T cell receptor (TCR) Vβ subset analyses

Total RNA was extracted from colon and spleen samples of 3 IL-10^-/-^, 3 SvEv WT, and 5 C57BL/6 mice and reverse transcribed to cDNA using random primers. Single-strand cDNA was used as template for PCR amplification of the complementarity-determining regions of the TCR using the Cβ primer and a set of 23 mouse Vβ primers, described previously [23]. Each Vβ-specific primer and Cβ primer were modified to contain the universal binding sequence at 5’ end for Illumina library construction with indexing primers to sequence the Vβ–Dβ junction region, to permit good coverage of CDR3 with Illumina pair-end reads. Final PCR products were gel purified and sequenced using Illumina Miseq sequencer (Illumina). Sequencing reads were quality filtered (Q value ≥20) and assigned germline Vβ/Jβ gene segments using MiXCR to identify TCR clonotypes for each Vβ region.

### Combination antiretroviral treatment (cART)

At 4 weeks of age, male wildtype (WT; n=17) and IL-10-/- mice (n=40) were randomly assigned to either a placebo or combined antiretroviral therapy (cART) for 12 weeks ± 1 week. The chosen cART regimen using repurposed HIV nucleoside/nucleotide reverse transcriptase inhibitors and protease inhibitors had been evaluated for activity in antagonizing MMTV *in vitro* and demonstrated antiviral activity *in vivo* corresponding with improvement of inflammatory disease in prior studies [21]. Animals were fed ad libitum with cART added to the drinking water, freshly prepared every second day for 12 weeks, to achieve a daily dose of 1 mg emtricitabine and 1.5 mg tenofovir disoproxil fumarate nucleoside/nucleotide reverse transcriptase inhibitors as well as 4 mg lopinavir boosted with 1 mg ritonavir protease inhibitors, as described previously [21]. Control treatment group mice received ground placebo tablets in their drinking water. Water consumption was monitored daily, and mouse weights measured weekly.

### Histological scoring

SvEv WT placebo (n=9), SvEv WT cART (n=8), IL-10^-/-^ placebo (n=21), and IL-10^-/-^ cART (n=19) mice were sacrificed after 12 weeks intervention. Colon weight and length were measured, and fresh colonic tissues were cut longitudinally and fixed in 10% phosphate buffered formalin, dehydrated overnight in 100% ethanol and prepared for paraffin embedding using an automated tissue processor. Formalin-fixed paraffin embedded (FFPE) tissue blocks were cut into 4 μm sections to be stained with H&E. Tissue images were assessed anonymously using a light microscope by a pathologist grading activity of each sample on a scale of 0 to 8 for enterocyte injury (0-3), epithelial hyperplasia (0-3), lymphocytes in lamina propria (0-2) and neutrophils in lamina propria (0-2), as previously described [18, 19]. Samples allotted a score of 0 to 2 are healthy with minimal damage to the gut lining and physiological inflammation, whereas samples with a score greater than 6 had marked diseased with increased inflammation and tissue damage.

### Evaluation of viral Load

Colons were collected from SvEv WT placebo (n=9), SvEv WT cART (n=8), IL-10^-/-^ placebo (n=21), and IL-10^-/-^ cART (n=19) mice after 12 weeks intervention for analysis of MMTV RNA levels. Initial quantification of MMTV RNA from untreated 16 to 18-week colon samples was performed using RT-qPCR to amplify target from 250 ng total colon RNA with primers complementary to MMTV Env normalized to Beta actin using the using the 2^−ΔCt^ **method,** as described [13].

The QuantiGene reagent system (Panomics/Affymetrix, Inc., Freemont, CA, USA) was used to measure response to cART, using methods previously described [24]. The QuantiGene is more precise than RT-PCR because the assay is capable of detecting 200 genome equivalents/sample and accurately quantifying RNA levels with > 400 copies/sample [24]. The probes were designed and synthesised by Panomics, using highly conserved regions in the *gag-pro-pol* genes from the MMTV genome (RefSeq NC_001503.1). QuantiGene RNA assay was performed on isolates, following manufacturer’s instructions. Briefly, RNA samples, including capture probes were added to a well with both blocking probes and capture extenders and incubated overnight. Samples were washed and amplified using a pre-amplifier solution. Samples were incubated for 1 hour, wash was repeated, and an amplifier solution was added. Label probes were added to each of the wells to allow chemiluminescent substrate binding to the label probes. Samples were measured using a luminometer and reported on an arbitrary, relative scale.

### MesoScale Discovery (MSD) Cytokine Assay

SvEv WT mice on placebo (n=9) or cART (n=8), and IL-10^-/-^ mice on placebo (n=19) or cART (n=21) were culled after 12 weeks intervention. Lysates from colonic tissue were analyzed using the Mesoscale Discovery (MSD) platform to identify cytokine secretion following manufacturer’s directions.

### Stool shotgun metagenomics library construction and bioinformatics

Total DNA was extracted from stool samples from SvEv WT mice on placebo (n=8) and cART (n=7), and IL-10^-/-^ mice on placebo (n=15) or cART (n=17), and cloned into Nextera XT libraries for Illumina sequencing [25]. Shotgun libraries sequences were quality-trimmed with fastq-mcf using a Q-score threshold of 24, a window size for trimming of 3 bp and a minimal read length of 120 bp. Quality-trimmed sequences were pseudo aligned with Kraken2 in paired-end mode [26] against a database including the NCBI RefSeq and the HMP databases with standard parameters and filtering hits that matched reference sequences with less that 10% of Kmers (-confidence 0.1). Furthermore, end1 and end2 reads were concatenated and aligned, after in silico translation, to the UniRef90 protein database using HUMAnN2 [27]. Kraken2 and HUMAnN2 alignment results were postprocessed using R scripts. Differential abundance analysis of taxa was modeled using a negative binomial distribution after scaling the data to account for sampling depth using the R package DESeq2 [28]. When compared against the control group, taxa were considered differentially abundant if the corrected P-value was smaller than 0.05. Representative plots were created with R scripts.

### Statistical Analysis

GraphPad Prism 9.0 was used for all statistical analyses. Data are expressed as means ± SEM with a minimum of 2 technical and 3 biological replicates. Differences between mean values were evaluated using the ANOVA with a post-test, student’s t-test, linear regression analysis to compare variables, and Fisher’s exact test to test categorical variables. TCR-VB analyses was conducted using a multiple unpaired t-test using a false discovery rate < 1% and two-stage step-up (Benjamini, Krieger, and Yekutieli).

## Results

### Characterization of MMTV infection in IL-10^-/-^ mice

While deriving evidence of MMTV infection in AMA producing mouse models with immune defects and spontaneous inflammatory disease [13], we observed increased expression of MMTV RNA in lymphoid tissues of IL-10^-/-^ mice, more so than the SvEv WT. In the present study, we directly addressed the question of whether the IL-10^-/-^ model expressed increased MMTV RNA in the colon using RT-qPCR and observed approximately a 10-fold increase in colon MMTV RNA in the IL-10^-/-^ as compared to the SvEv WT **(Figure 1A).** As neonatal mice make poor humoral responses to MMTV infection [17], we sought evidence of cellular immune responses to MMTV instead. For these studies we examined production of the proinflammatory cytokine IFN-γ in both the SvEv WT and the IL-10^-/-^ mice by stimulating splenocytes with overlapping MMTV Gag peptides **(Supplementary Table 1**). We found the IL-10^-/-^ mice demonstrated a slight increase in IFN-γ response production at baseline, and therefore, the stimulation studies were normalized by subtracting the background cytokine levels **(Figure 1B).** Following stimulation with MMTV Gag peptides, splenocytes from IL-10^-/-^secreted increased levels of IFN-γ as compared to the SvEv WT (0.97 vs. 0.10 pg/ml, p=0.031), reflecting augmented proinflammatory responses to MMTV.

**Figure 1.**
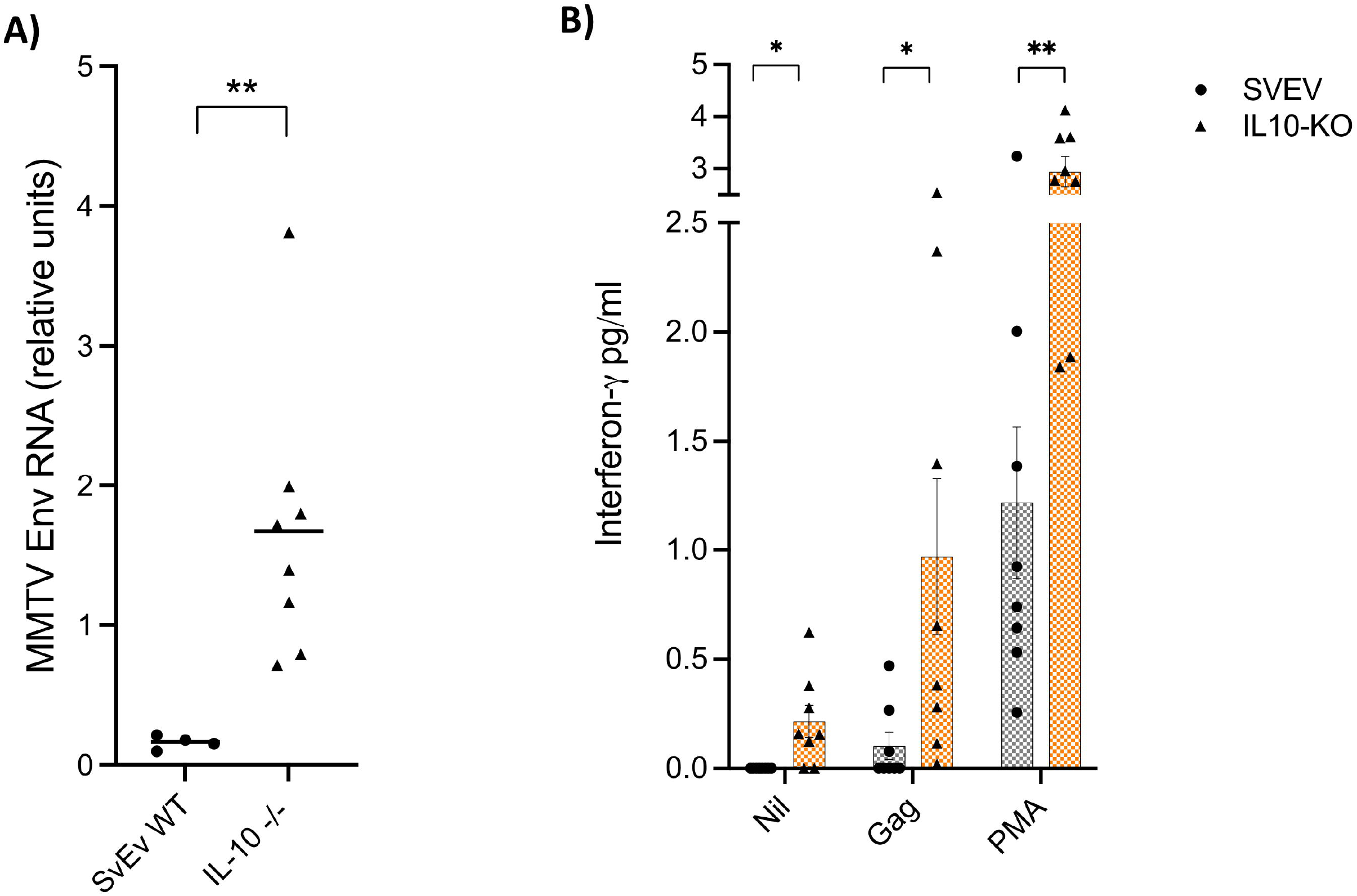
Increased viral load and immune responses to MMTV in IL-10^-/-^ and SvEv WT mouse. **(A)** RT-PCR of colon RNA shows increased relative MMTV copy number in IL-10^-/-^ vs. SvEv colon (relative units; **p=0.004, Mann Whitney test). (B) Interferon-γ release assay showing increased production in the IL-10^-/-^ vs. SvEv splenocytes without any stimulus (Nil) and following stimulation with either MMTV Gag peptides or PMA/ionomycin [Mean±SEM, Gag and PMA levels normalized to background production; *p=0.05, **p=0.002, unpaired t-test].

As exogenous MMTV is milk borne, evidence of enteral infection can be found in weanling pup stomachs [14]. Accordingly, we made viral preparations from stomach lactoserum using the cell pellets from the stomach as controls and cloned total RNA into next generation sequencing libraries using a capture method to enrich the betaretrovirus sequences. By comparing read abundance in the Illumina sequencing libraries, more than a 10-fold enrichment of MMTV reads was observed in the IL-10^-/-^ compared to the SvEv WT lactoserum libraries (**Table 1**). Calculations of genome equivalents derived by RT-qPCR prior to the capture enrichment revealed a viral abundance of approximately 3.0 x 10^8^ in the IL-10^-/-^ lactoserum consistent with milk borne infection [14]. The increased representation of MMTV reads in lactoserum versus cell pellet libraries is also consistent with enrichment of viral particles in the mouse milk (**Table 1**).

**Table 1.** Estimates of relative MMTV copy number calculated by number of MMTV reads per illumina library and RT-PCR copy number in lactoserum RNA.

|  | *MMTV reads*<br>*per library* | *MMTV reads %*<br>*per library* | *Average read depth*<br>*per MMTV genome* | *RT-PCR MMTV*<br>*copy number\** |
| --- | --- | --- | --- | --- |
| **SvEv stomach pellet** | 174,491 | 1.53% | 3,548 x |  |
| **SvEv lactosera** | 220,375 | 13.15% | 5,132 x | 1.2 x 10<sup>6</sup> |
| **IL-10 stomach pellet** | 717,773 | 6.96% | 15,270 x |  |
| **IL-10 lactosera** | 2,886,869 | 51.33% | 67,017 x | 3.0 x 10<sup>8</sup> |

\* RT-PCR estimate of copy number x (amount of total RNA extracted from lactosera viral preparation/amount of total RNA amount used for RT-PCR reaction).

Analysis of the mouse milk viral prep showed that over half of the Illumina reads in the IL-10^-/-^ lactoserum library aligned with MMTV to provide a 99.40% coverage of the viral genome (**Table 1, Supplemental Figure 1**). Following assembly, blastn searches showed that the nine MMTV contigs were mainly related to the endogenous *Mtv-1* loci and the exogenous MMTV(HeJ) virus from C3H mice, which is a genetic recombinant of the endogenous *Mtv1* provirus and exogenous MMTV(C3H) variant [22]. The two longest contigs shared 96.4% to 97.2% identity with the MMTV(HeJ) variant, whereas the shorter contigs shared identity to *env* and *sag* genes from *Mtv1* **(Table 2).** The MMTV genome derived from the IL-10^-/-^ lactoserum showed a degree of variability in the longest two contigs with 98.4% identity with each other **(Table 2;** scaffolds 1 and 3). The highest region of variability is the 3’ region of the *sag* gene, whereas different exogeneous MMTV strains demonstrate limited variability as compared to other retroviruses, such as HIV, due to the high fidelity of the MMTV Pol protein that minimises error during the reverse transcription process [29].

**Table 2.**
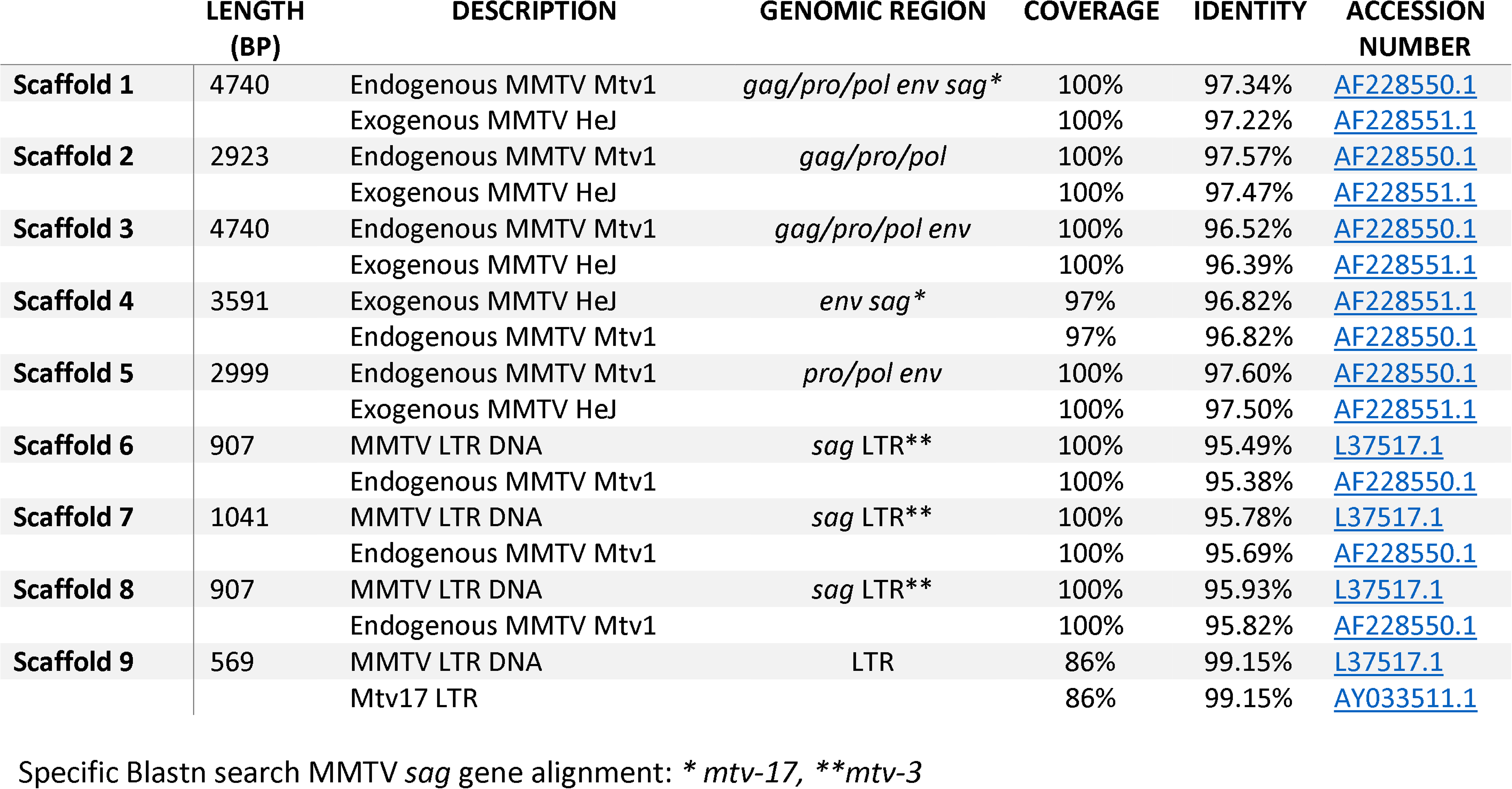
Blastn analyses of contigs derived from IL-10^-/-^ lactoserum showing the top two significant alignments.

|  | **LENGTH**<br>**(BP)** | **DESCRIPTION** | **GENOMIC REGION** | **COVERAGE** | **IDENTITY** | **ACCESSION**<br>**NUMBER** |
| --- | --- | --- | --- | --- | --- | --- |
| **Scaffold 1** | 4740 | Endogenous MMTV Mtv1 | *gag/pro/pol env sag\** | 100% | 97.34% | AF228550.1 |
|  |  | Exogenous MMTV HeJ |  | 100% | 97.22% | AF228551.1 |
| **Scaffold 2** | 2923 | Endogenous MMTV Mtv1 | *gag/pro/pol* | 100% | 97.57% | AF228550.1 |
|  |  | Exogenous MMTV HeJ |  | 100% | 97.47% | AF228551.1 |
| **Scaffold 3** | 4740 | Endogenous MMTV Mtv1 | *gag/pro/pol env* | 100% | 96.52% | AF228550.1 |
|  |  | Exogenous MMTV HeJ |  | 100% | 96.39% | AF228551.1 |
| **Scaffold 4** | 3591 | Exogenous MMTV HeJ | *env sag\** | 97% | 96.82% | AF228551.1 |
|  |  | Endogenous MMTV Mtv1 |  | 97% | 96.82% | AF228550.1 |
| **Scaffold 5** | 2999 | Endogenous MMTV Mtv1 | *pro/pol env* | 100% | 97.60% | AF228550.1 |
|  |  | Exogenous MMTV HeJ |  | 100% | 97.50% | AF228551.1 |
| **Scaffold 6** | 907 | MMTV LTR DNA | *sag* LTR** | 100% | 95.49% | L37517.1 |
|  |  | Endogenous MMTV Mtv1 |  | 100% | 95.38% | AF228550.1 |
| **Scaffold 7** | 1041 | MMTV LTR DNA | *sag* LTR** | 100% | 95.78% | L37517.1 |
|  |  | Endogenous MMTV Mtv1 |  | 100% | 95.69% | AF228550.1 |
| **Scaffold 8** | 907 | MMTV LTR DNA | *sag* LTR** | 100% | 95.93% | L37517.1 |
|  |  | Endogenous MMTV Mtv1 |  | 100% | 95.82% | AF228550.1 |
| **Scaffold 9** | 569 | MMTV LTR DNA | LTR | 86% | 99.15% | L37517.1 |
|  |  | Mtv17 LTR |  | 86% | 99.15% | AY033511.1 |

Specific Blastn search MMTV *sag* gene alignment: \* *mtv-17*, \*\**mtv-3*

Active MMTV infection in mice can be demonstrated by evaluating superantigen activity and observing differences in the cognate TCR-Vβ subsets that react with the specific vSAG [14]. The IL-10^-/-^ model has a mixed genetic background of several mouse strains including the C57BL/6 and sub strains of 129, each with different and partially characterized endogenous *mtv* loci [30]. For example, the C57BL/6 mouse has three full length endogenous genomes, *mtv-8, mtv-9* and *mtv-17*, and sub strains of 129 mice contain combinations of *mtv-1, mtv-3, mtv-8, mtv-9, mtv-11, mtv-13*, and *mtv-17* [14–16]. As variable *sag* sequences were observed in the IL-10^-/-^ mouse milk **(Table 2),** we cloned and sequenced MMTV *sag* directly from spleen RNA to determine the predominant vSAG triggering changes in TCR-Vβ subsets. Eleven of 12 clones were derived from *mtv-9 sag* encoding the vSAG9 **(Figure 2A)** that preferentially binds and expands TCR-Vβ5, TCR-Vβ11, and TCR-Vβ12 lymphocytes [14]. We then evaluated the TCR-Vβ subset distribution in the spleen and colon **(Supplemental Figure 2)** and found that TCR-Vβ5, and TCR-Vβ12 were expressed in sufficient quantity for analysis, but TCR-Vβ11 constituted less than 0.5% of the population. Significant differences were observed in the colon that were not observed in the spleen **(Supplemental Figure 2).** Consistent with vSAG9 induced activity, the percentage of TCR-Vβ12 was significantly increased (17.14 vs. 6.45, q=0.012) and a trend was observed for increased TCR-Vβ5 (IL-10^-/-^ vs. SvEv, 0.63% vs.0.21%, q=0.067) in the IL-10^-/-^ versus the SvEv WT colon **(Figure 2B).**

**Figure 2.**
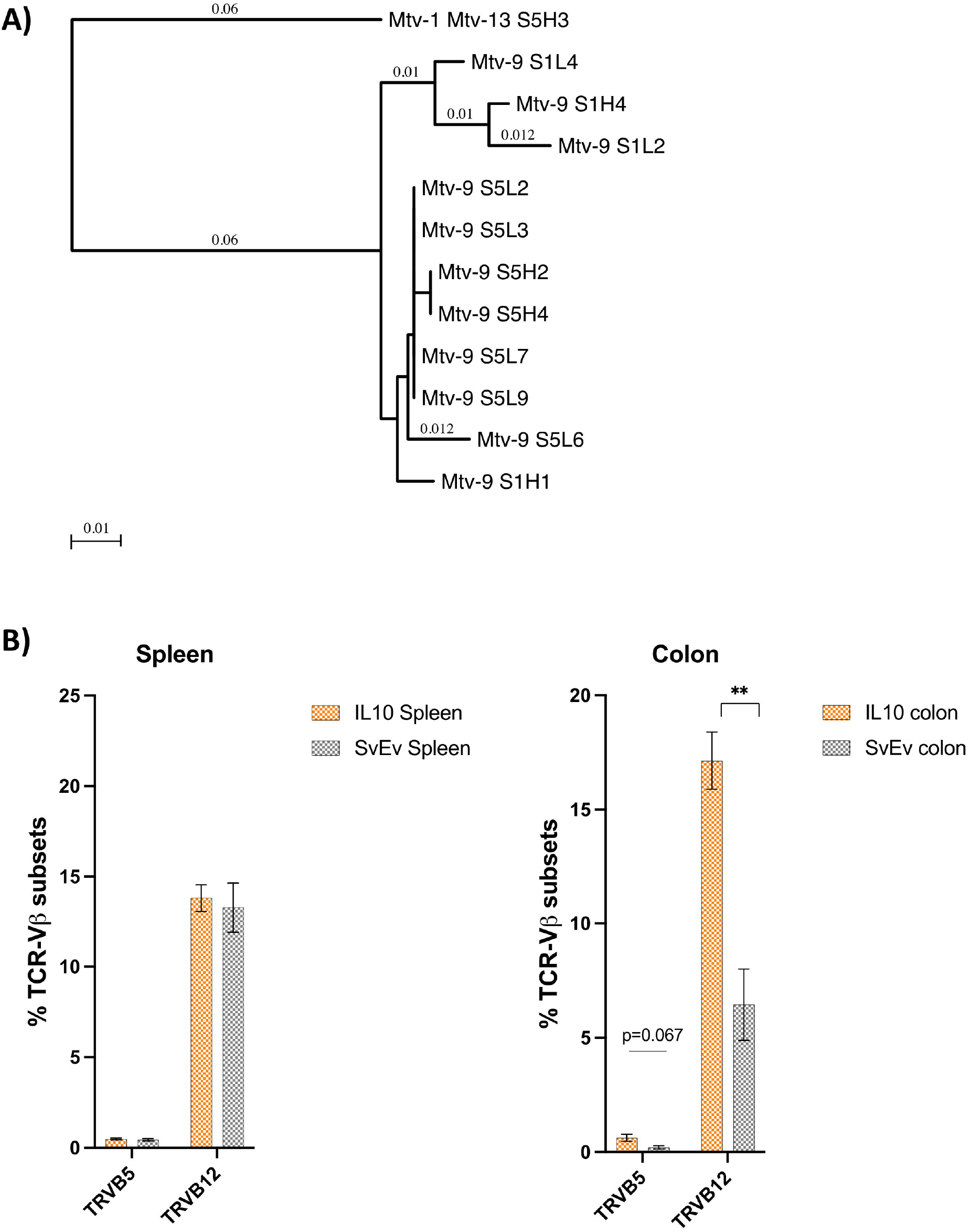
MMTV superantigen and cognate TCR-Vβ differences in IL-10-/- vs. SvEv WT colon. **(A)** Phylogram of sag clones showing a predominance (92%) of mtv-9 that activates TCR-Vβ5 and TCR-Vβ12 (Phylogenetic tree made by MacVector 18.5: Neighbor Joining; Best Tree; tie breaking = random; Distance: uncorrected (“p”); gaps distributed proportionally). (B) TCR-Vβ5 and TCR-Vβ12 subset distribution of read count assessed by Illumina sequencing in spleen and colon showed no significant differences between IL-10-/- vs. SvEv in the spleen, whereas IL-10-/- colon had more than 2-fold % increase in TCR-Vβ5 and TCR-Vβ12 [Mean±SEM, ** p=0.006, Multiple unpaired t-test, Benjamini, Kreiger, and Yekutieli two stage set up].

### Combination ART reduces the severity of colitis in IL-10^-/-^

To address the hypothesis that MMTV plays a role in the development of inflammation in the IL-10^-/-^ mouse model, we employed a cART regimen previously shown to ameliorate MMTV cholangitis in the NOD.c3c4 autoimmune biliary disease model of primary biliary cholangitis [21]. The cART was composed of combination nucleoside reverse transcriptase inhibitors and a boosted protease inhibitor added to the drinking water of IL-10^-/-^ and SvEv WT mice **(Figure 3A).** The cART and placebo were commenced at 4 weeks of age and following this, mice were assessed for body weight every 2 weeks. The cART treatment did not alter growth in either WT or IL-10 ^-/-^ mice **(Figure 3B).** The weight gain over time was reduced in the IL-10 ^-/-^ pups (n=10) versus WT (n=4) receiving cART and similarly in the IL-10^-/-^ (n=9) versus WT (n=4) receiving placebo (p<0.0001 for both comparisons). As both WT and IL-10 ^-/-^ a similar amount of food and drink and the drop off in weight corresponds with the development of colitis, we attribute the weight changes to catabolism from the inflammatory bowel disease.

**Figure 3:**
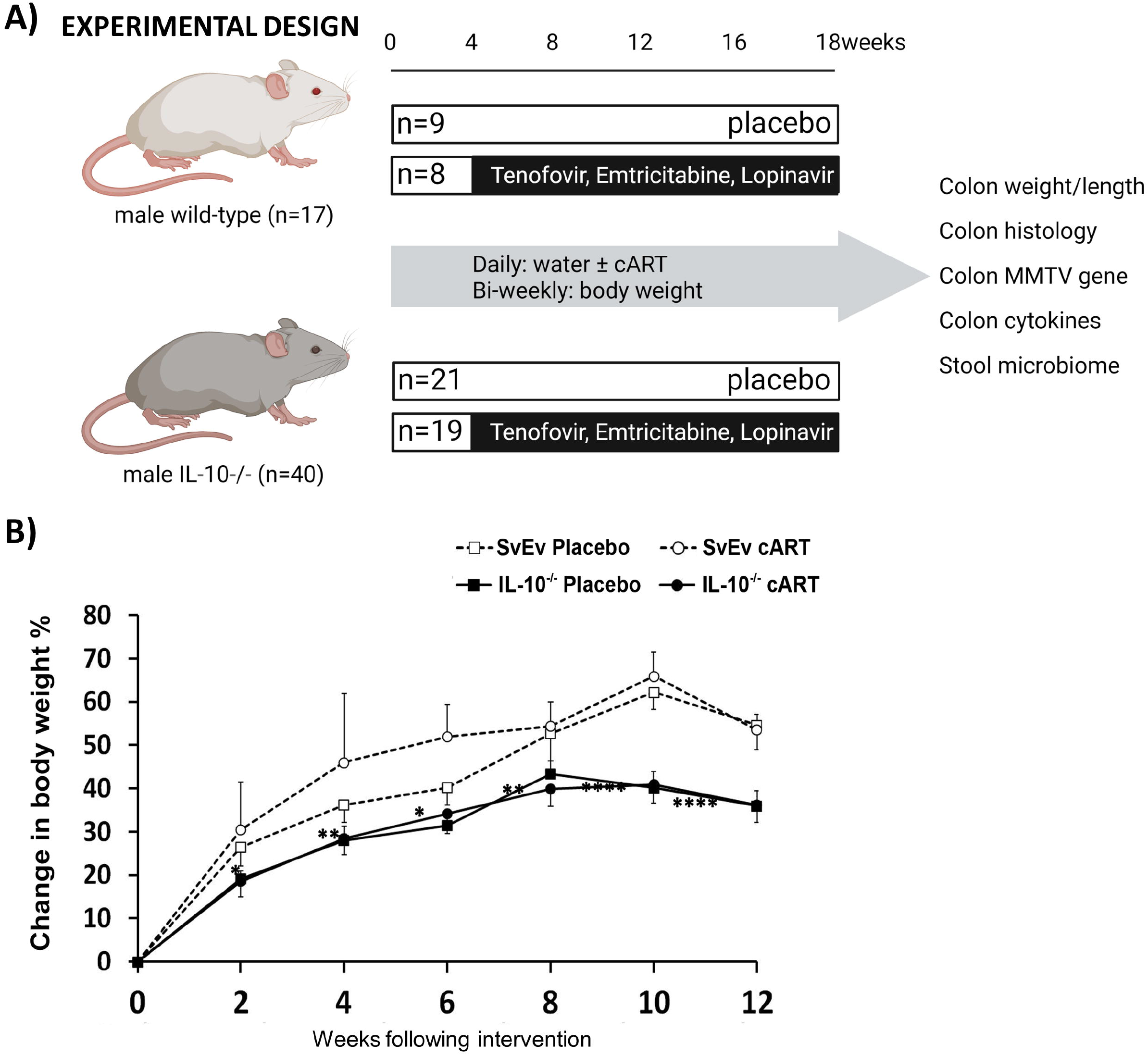
Antiretroviral therapy does not impact on growth changes in IL-10-/- mice. **(A)** SvEv WT and IL-10-/- mice commenced intervention aged 4 weeks with control (placebo) or cART for 12 weeks ± 1 week with daily nucleoside reverse transcriptase inhibitors (1 mg emtricitabine and 1.5 mg tenofovir disoproxil fumarate) and protease inhibitors (4 mg lopinavir boosted with 1 mg ritonavir) in drinking water. Body weight was continuously measured. Interventions occurred at 12 weeks at which times histology tissue scores, mucosal cytokine production, and viral assessments were performed. (B) Mouse weight, expressed as a percentage change compared to week 0, was measured every 2 weeks over a total of 12 weeks [Mean±SEM, ****p<0.0001, **p<0.01, *p<0.05].

As anticipated, IL-10 ^-/-^ mice had greater histological damage in the colon as compared to the SvEv WT animals (p<0.001, **Figure 4A, 4B).** The cART was associated with reduced histological scores in the IL-10^-/-^ mice as compared to those on placebo (p<0.01), which was associated with a significant reduction in MMTV viral load **(Figure 4C).** Whereas WT treated with cART had minimal change versus placebo in histology or MMTV RNA expression, which likely reflected endogenous retrovirus expression that would not be responsive to antiviral therapy **(Figure 4B, 4C).** Colon weight to length ratio was significantly increased in IL-10 ^-/-^ mice compared to SvEv WT (p<0.002). Also, the colon weight to length ratio in IL-10^-/-^ mice on cART showed a trend for reduction compared to IL-10^-/-^ on placebo **(Figure 4D).**

**Figure 4:**
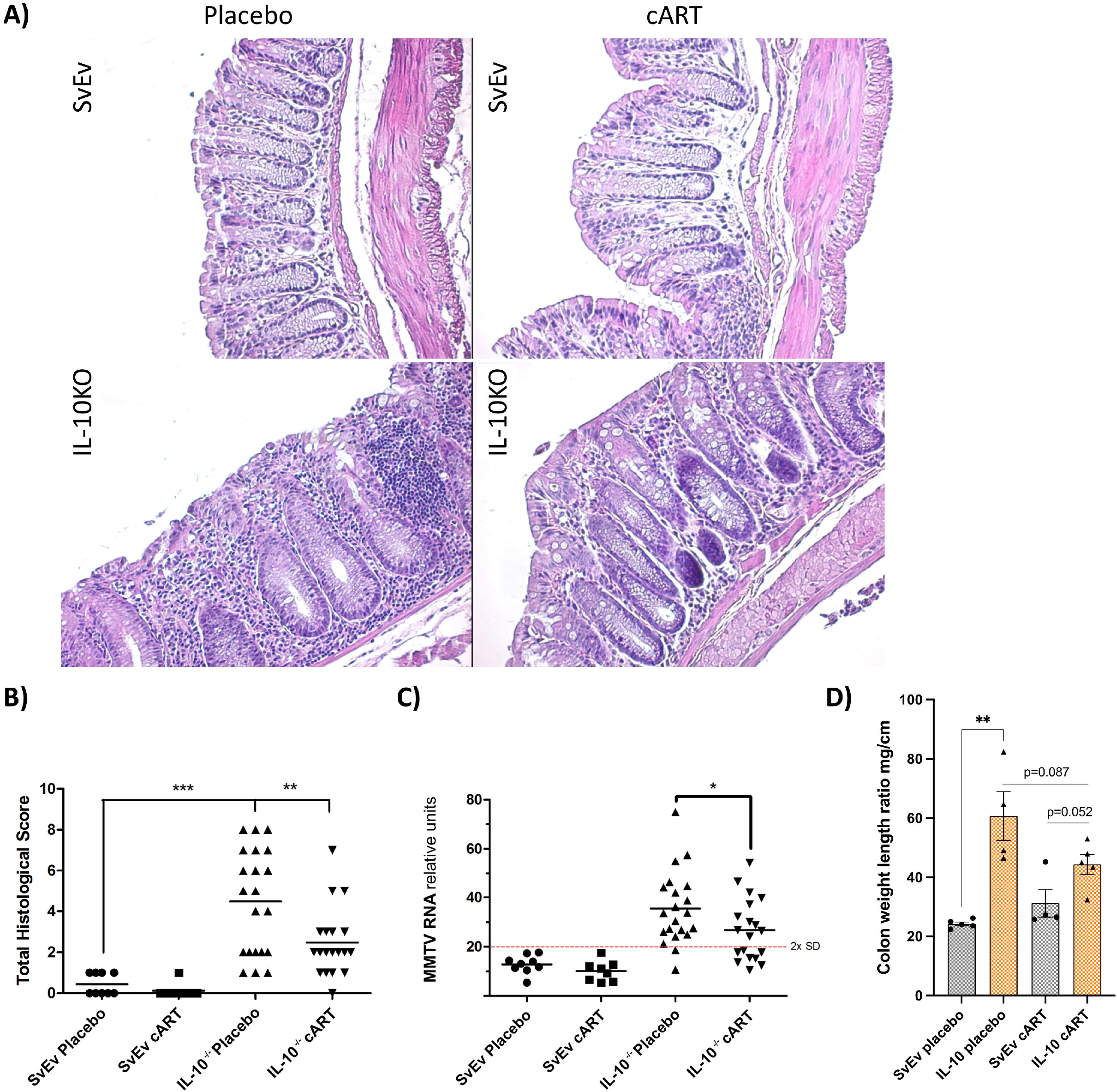
cART reduces histological score and MMTV viral load in colon tissues of IL-10-/- mice. **(A)** Hematoxylin and Eosin stain of colon tissues shows marked reduction in inflammation in the IL-10-/- sample following cART to levels observed in the SvEv wildtype. (B) Histological scoring of colon tissue samples [***p<0.001, **p<0.01]. **(C)** Colon MMTV RNA load was reduced 25% in IL-10-/- mice receiving cART vs. placebo; 42% of IL-10-/- mice receiving cART developed background MMTV levels versus 10% of those on placebo [*p<0.028, Fishers exact test. Cutoff = 2xSD of background SvEv WT colon samples]. **(D)** Colon weight to length ratio used as a surrogate of gut inflammation was evaluated after 12 weeks of treatment [Mean±SEM, **p=0.0015].

To evaluate the role of viral infection in generating colitis in the IL-10^-/-^ mouse model, we measured the levels of MMTV RNA in the colon using QuantiGene hybridization technology rather than RT-qPCR because of the improved accuracy of the former with lower levels of MMTV RNA [24]. Of note, these levels represent both the presence of exogenous MMTV viral RNA as well as the expression of endogenous viral mRNA. A cutoff was established using background levels MMTV RNA in SvEv WT colon samples **(Figure 4D)** and as a result, all the WT mice were found to have colon MMTV RNA levels below this level as compared to 25% of the IL-10^-/-^ (Fisher’s exact test, p<0.0001). Little difference was observed in the WT mouse colon MMTV RNA levels with cART, whereas the mean levels were reduced 25% in IL-10^-/-^ mice receiving cART vs placebo **(Figure 4D).** Following cART, 42% of IL-10^-/-^ colon samples were found to have background levels of MMTV RNA as compared 10% of the mice of placebo (Fishers exact test, p<0.028). Also, a trend was observed for a correlation of viral load and histology score (Spearmen r 0.32, p=0.082, **Supplementary Figure 3)**

To gain a more in depth understanding of the effects of cART on inflammation in the IL-10^-/-^ mouse model, we examined presence of a series of IBD-associated cytokines in colonic tissues. As expected, the levels of the pro-inflammatory markers, TNFα, KC GRO, IFNγ, IL-6, IL-1β, IL-12p70, IL-4, and IL-5 were generally elevated in IL-10^-/-^ mice compared to SvEv WT **(Figure 5, Supplementary Figure 4).** At the end of study, the IL-10^-/-^ mice on cART experienced a significant reduction in the levels of TNFα, KC GRO and IL-6 in colon tissue lysates compared to placebo (p<0.05, **Figure 5**).

**Figure 5:**
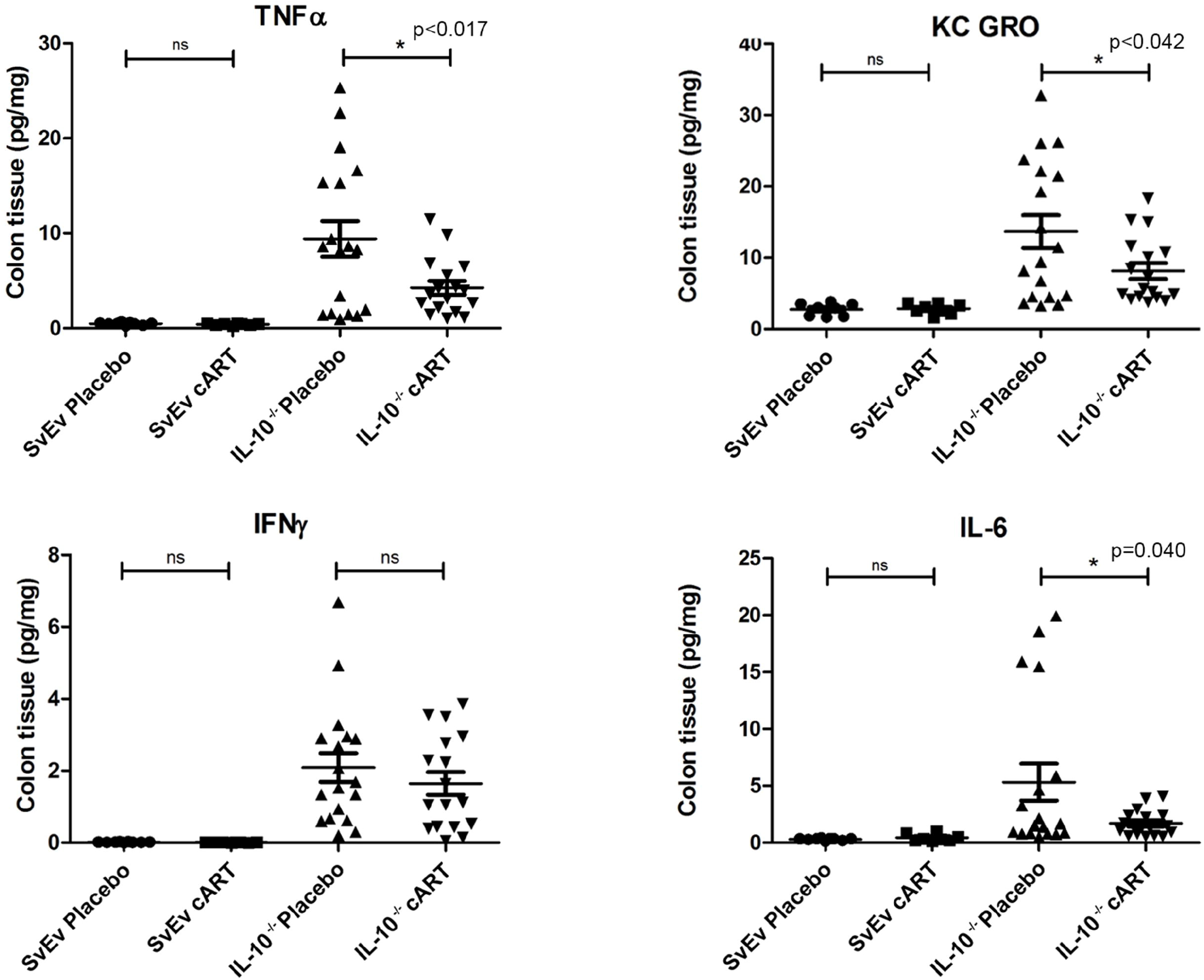
cART impacts on colon pro-inflammatory cytokines. **Colon** extracts were analyzed for pro-inflammatory cytokines associated with IBD. Levels of TNFa, KC GRO, IFNg and IL-6 were were elevated in IL-10-/- mice versus WT mice and the cytokines TNFa, KC GRO and IL-6 were altered in the IL-10-/- mice on cART [Mean±SEM, *p<0.05].

### cART causes shift in microbial abundance profiles of IL-10^-/-^mice

As the virome plays a key role in homeostasis of the microbiome community [4], we sought to examine the effect of cART treatment on the gut microbiome of IL-10^-/-^ mice. As seen in Figure 6, cART treatment induced chnages in the microbial composition in the IL10^-/-^ mice. For example, an increased abundance in *Absiella inocuum, Absiella*, and *Blautia* was seen in the IL10^-/-^ mice treated with cART versus placebo, and a decreased abundance of *Lactobacillus johnsonii, Lactobacillus taiwanensis*, and *Staphylococcus xylosus* in IL10^-/-^ mice treated with cART compared to placebo **(Figure 6A).** In contrast, there were no significant changes in abundance of microbial taxa in SvEv WT mice treated with ART compared to placebo **(Figure 6B).**

**Figure 6:**
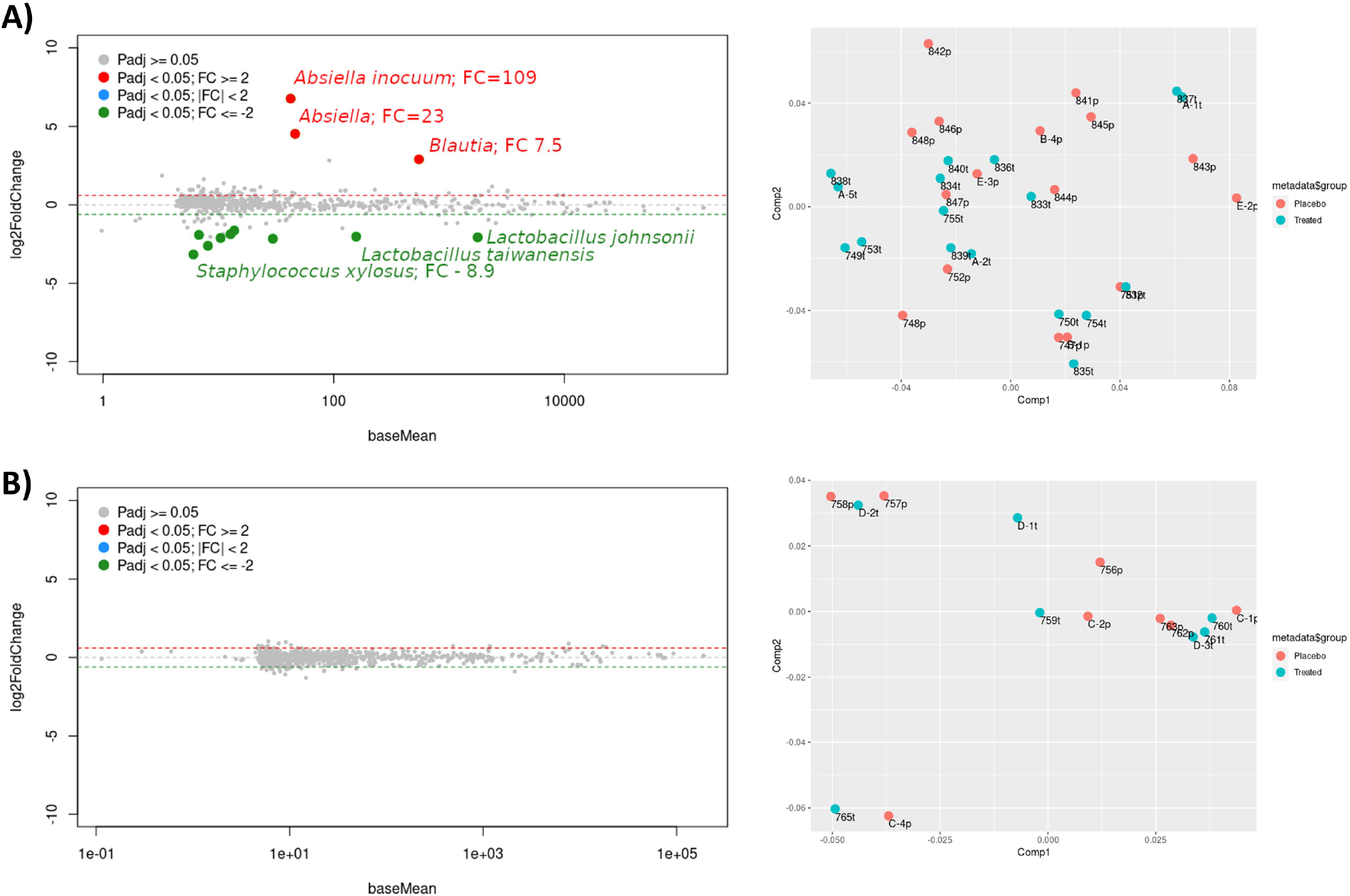
Differences in taxa abundance in response to cART in IL-10-/- vs. SvEv WT. Comparisons between animals in the placebo and cART groups show that the abundance of some taxa were significantly different in **(A)** the colitis model IL-10-/- but not **(B)** the SvEv wildtype. [Left panels show differential abundance analyses with the package DESeq2; red and green dots represent taxa with increased or decreased abundance, respectively, in the ART group when compared against placebo and only taxa with a fold change difference of 2 or more are shown. Right panels represent the first and second components of a principal coordinate analysis using Bray-Curtis distances; number of animals in the placebo and antiviral therapy groups, respectively, were IL10-/-: n=19 and n=21; SvEv: n=9 and n=8].

## Discussion

The etiology of IBD is complex, involving genetics, a dysregulated immune system, environmental factors, and the interaction of the virome within the microbiome [1, 3, 4, 31]. Herein, we characterized a betaretrovirus infection in the IL-10 ^-/-^ model of colitis, which is endemic in laboratory mice, replicates in the mucosal lymphocytes using a viral superantigen mechanism, and subverts immune responses by triggering the production of IL-10 [14, 17]. We report an increased viral load of MMTV in breast milk and colon of the IL-10^-/-^ mouse, proinflammatory cellular responses to MMTV and viral superantigen activity in the colon to support a hypothesis that MMTV infection is linked with the augmented inflammatory responses in the IL-10^-/-^ model of colitis. We found a correlation between MMTV and inflammation in the IL-10^-/-^ colon, as demonstrated by an increased viral load, histopathology score and pro-inflammatory cytokines versus the WT and then using antiviral treatment with known activity against MMTV [21], we demonstrate a definitive response to cART with decreased viral load, histology score, and inflammatory cytokines, linked with changes in the microbiome.

Several features of the MMTV infection may combine with IL-10 deficiency to trigger colitis in this model. IL-10 plays an important role in tolerizing neonatal mice to viral infection as part of an important interplay of MMTV with commensal microbiota in the gut. Following milk borne infection, MMTV bound bacterial-LPS engages tolllike receptor 4 on M cells or B cells in the Peyer’s patches, which in turn triggers production of IL-10 in an IL-6 dependant fashion. The two important features of this process are that MMTV requires commensal gut microbes for productive infection, and that production of IL-10 limits antiviral responses by preventing the formation of neutralizing antibodies (and possibly cellular immunity as well) [17]. Relevant to the IL-10^-/-^ model, colitis is abrogated in germ free conditions and ameliorated by changing the balance of intestinal flora, which may be related to the requirement of MMTV to use bacterial LPS to cross the mucosal barrier and gain access to the Peyer’s patches [18, 19].

Infection by the oral route also requires MMTV replication in proliferating CD4^+^ lymphocytes using a vSAG dependant mechanism in the gastrointestinal tract [14, 17]. Indeed, providing evidence of ongoing superantigen activity with cognate TCR-Vβ expansion is used to demonstrate active MMTV infection in mouse models [14, 17]. The superantigen activity is both dynamic and complex because vSAG reactive TCR-Vβ CD4^+^ lymphocytes are initially activated and then depleted, and this may occur either during embryonal development or during replicative infection, and the lymphocyte responses vary in a tissue-dependant and mouse strain-dependant fashion [14]. In this study, identification of the relevant vSAG was not straight forward because different MMTV *sag* genes were identified in the IL10^-/-^ lactoserum contigs **(Table 2;** *mtv-3* specific for TCR-Vβ3, TCR-Vβ5 and TCR-Vβ17 and *mtv-17* of unknown specificity [14]). Ultimately the vSAG present in infected tissue dictates specificity, and a vSAG9 effect was observed with more than a 2-fold increase of the cognate TCR-Vβ5 and TCR-Vβ12 subsets in the IL10^-/-^ colon.

Other features consistent with MMTV infection and vSAG activity have been previously observed in the IL-10^-/-^ model. For example, a marked expansion of the colon CD4+ T-cell population is found in the IL-10^-/-^ mouse leading to a reversal of the CD4^+^/CD8^+^ T-cell ratio found in WT mice [18]. The vSAG activation of CD4+ lymphocytes leads to a progressive increase cognate TCR-Vβ Foxp3+ regulatory T cells (Treg) in Peyer’s patches and these MMTV infected Treg have the ability to preferentially suppress the proliferative response of T cells *ex vivo* in wildtype mice [32]. Whereas transfer of purified CD4+ lymphocytes from IL-10^-/-^ mice to immunodeficient RAG mice results in the development of chronic colitis, with the CD4+ population as the predominant culprit [33]. Under these circumstances, the colitis in the RAG mice is presumably linked with both the passage of MMTV infection and lack of IL-10 activity.

Our findings may be clinically relevant as vSAG activity has been suggested as a potential mechanism in CD pathogenesis with TCR-Vβ skewing and evidence of enrichment of CD4^+^ lymphocytes in inflamed mucosa [34]. However, vSAG activity has been hypothesized as pathogenic mechanism in multiple human autoimmune and idiopathic inflammatory diseases without further identification of a viral superantigen to better delineate cognate TCR-Vβ skewing. With regard to the predominance of CD4^+^ lymphocytes in Crohn disease, the observation that human immunodeficiency virus is linked with decreased disease severity in patients with IBD has seeded the idea of the “remission theory”, which suggests that the lower CD4+ counts translate to a diminished disease process [35]. This idea too requires further evaluation and does not consider that the use of cART may also impact on the pathogenesis of IBD as demonstrated herein with the IL-10^-/-^ model [35].

The “virus + susceptibility gene” model of colitis is the IL10^-/-^ mouse is confounded somewhat because part of the “susceptibility” is related to the development of MMTV infection. Indeed, the parental strains do not manifest phenotypic features of MMTV infection but infection emerges in the absence of IL-10 [36]. A similar process has been reported in other models following genetic manipulation to create immune deficiency. For example, the C57BL/6 mouse does not express infectious gamma-retroviruses in health, but following genetic disruption of antibody responses, the model develops spontaneous murine leukemia virus infection from endogenous loci and this is accompanied by the development of retrovirus induced lymphoma [12]. As discussed, C57BL/6 encode several full-length endogenous MMTV proviral genomes that are heavily methylated to silence viral RNA expression and encode multiple mutations to prevent production of viral proteins [13]. Whereas other lines, such as C3H, are less resistant to MMTV infection and produce MMTV particles in milk that leads to the development of breast cancer in offspring [14]. The effects of MMTV in 129/SvEv are largely unknown; sub strains are frequently used for breeding due to the availability of embryonal stem cells and in general, these mice are not prone to development of cancers linked with MMTV. It is possible that the SvEv WT may exhibit low level MMTV production as shown by the enrichment of MMTV reads in the lactoserum versus stomach cell pellet library (**Table 1**), but this requires further evaluation as the SvEv WT shows no appreciable phenotype of infection. Another consideration is that the IL-10 knock out “susceptibility gene” is only linked with colitis in specific lines of inbred mice, such as 129/SvEv and C3H/HeJBir, whereas the C57BL/6 are relatively resistant to developing intestinal disease, suggesting that other genetic modifiers play a role in the process [36]. Therefore, it is tempting to speculate that a portion of the genetic background contributing to the development of colitis may include the susceptibility to MMTV or other viral infection as well.

Other considerations for any IBD model are the global versus specific effects mediated by the “susceptibility gene” lesion. For example, the T cell TGF-β receptor II dominant-negative mouse (C57BL/6/C3H background) and the interleukin-2 receptor α knock out mouse (C57BL/6 background) develop diffuse multi-organ inflammatory disease associated with IBD and autoimmune biliary disease; and both models exhibit markedly elevated levels of MMTV RNA and proteins in lymphoid tissue and inflamed organs [13, 37, 38]. In these models, one can argue that the diffuse disease process is likely related to the global immunodeficiency that permits emergence of endogenous retroviruses in general (even on the C57BL/6 background), and the potential for infection with other exogenous pathogens. In contrast, the IL-10 deficient models tend to develop disease restricted to the gut, which is likely due to the predominant role IL-10 plays in mediating intestinal homeostasis by down-regulating Th1 responses, NK cell activity, dendritic cell antigen presentation, and pro-inflammatory cytokine production [18].

Interestingly, while a dysbiotic microbiota (including bacteria, fungi, and viral species) is a hallmark of IBD in humans and is seen in IL-10^-/-^ mice, we only observed small effects of cART therapy on microbial composition. Significantly, cART in the IL10^-/-^ mice resulted in an increased abundance of the probiotic commensal microbe, *Blautia*, which is associated with remission in IBD patients [39]. Meanwhile, *Lactobacillus* species which have previously been associated with disease flare in IBD patients [40], were decreased following cART in the IL10^-/-^ mice. These findings suggest that MMTV may play a minor role in cross communication between microbiome viruses and bacteria [17] although in our model, cART did not significantly affect bacterial abundance patterns in IL10^-/-^ mice. Rather than microbial abundance, MMTV may instead play a larger role in microbial functions.

A caveat of these studies was the use of male mice only, and the indirect method employed to reduce MMTV replication. To reduce heterogeneity of responses, male mice were used for the study because MMTV is activated by female hormone response elements and viral replication is triggered more so during female puberty. While female IL-10 mice may develop worse colitis on occasion, this is not a consistent observation. Also, the combination ART used to treat MMTV was repurposed from treating the lentivirus, HIV, and not specific for betaretrovirus infection. Previously, we demonstrated that each of the components of this cART regimen inhibits MMTV replication and together improved MMTV induced cholangitis [21]. Nevertheless, we cannot rule out inhibition of other retroviruses nor off target effects in modulating the model even though MMTV levels were reduced with the cART.

In conclusion, in this study we have demonstrated that MMTV virus is present in the IL-10^-/-^ mouse model of colitis and contributes to severity of inflammation. These results suggest that MMTV may contribute to colitis in this model, although we recognize that we did not investigate other retroviruses that could also be involved. The outcomes of this study may also help us better understand the interplay between commensal bacteria and viruses leading to more effective therapies and prevention techniques for mucosal transmission of viruses. Our studies also provide an impetus to continue investigating the role of viruses in the etiology of IBD.

## DECLARATIONS

### Ethical approval

The protocol was approved by the University of Alberta Health Research Ethics Board (Study ID. AUP00000294).

### Consent for publication

All authors have read and approved the final version of the manuscript for publication.

### Availability of data and materials

All data are included in the manuscript.

### Authors’ contributions

KM, BR, and AM conceived and designed the experiments. NH, HS, NS and WW performed the experiments. HA HK, HS, NS, MR and WW were responsible for the figure preparation and statistical analyses. HA and AM drafted the manuscript. All authors edited and approved the final version of the manuscript.

## Acknowledgements

We would like to acknowledge^1^ CEGIIR and The Applied Genomic Core, University of Alberta, for providing material support including Chelsea McDougall for assistance with animal maintenance and collections, Juan Jovel for sequencing analysis, and Harsh Thaker for experimental support.

---

**Supplementary Figure 1:**
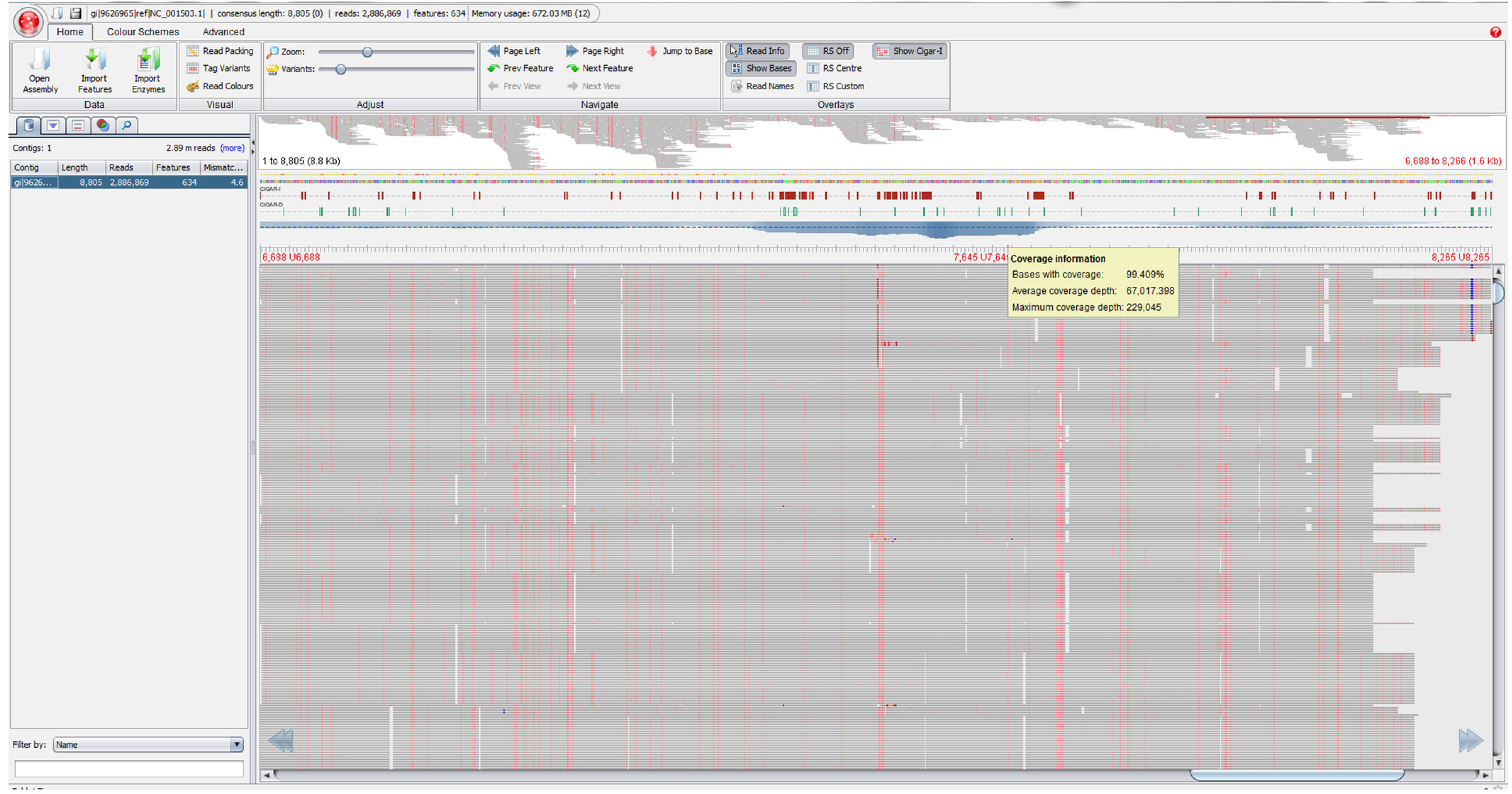
Alignment of Illumina reads from IL-10-/- lactoserum library to MMTV genome showing a screen capture with alignment of 2,886,869 million Illumina reads along MMTV reference genome with variance in sequence marked in red, to provide 99.40% coverage of the MMTV genome and an average read depth of 67,017-fold.

**Supplementary Figure 2:**
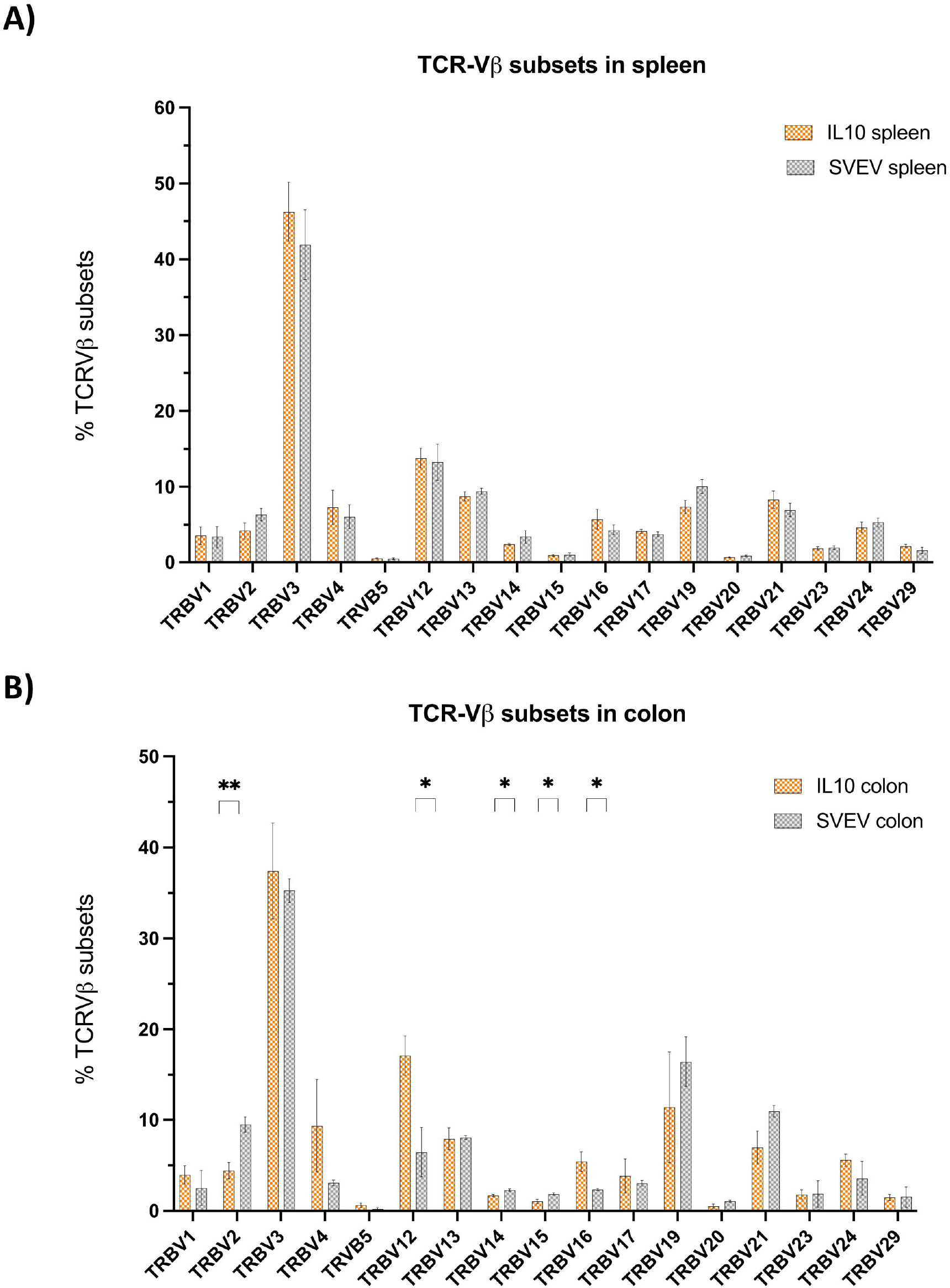
T cell receptor (TCR)-Vβ subset distribution in spleen and colon of IL-10-/- vs. SvEv mice assessed by Illumina sequencing. (A) No significant differences were observed between IL-10-/- vs SvEv in the spleen. (B) the IL-10-/- colon had increased TCR-Vβ12 and TCR-Vβ16 subsets with diminished TCR-Vβ2, TCR-Vβ14 and TCR-Vβ15 expression. [Mean±SEM, TCR-Vβ subsets with percent less < 0.5% removed from the analyses. * p<0.01, ** p=0.002, Multiple unpaired t-test, Benjamini, Kreiger, and Yekutieli two stage set up, q value < 0.1].

**Supplementary Figure 3:**
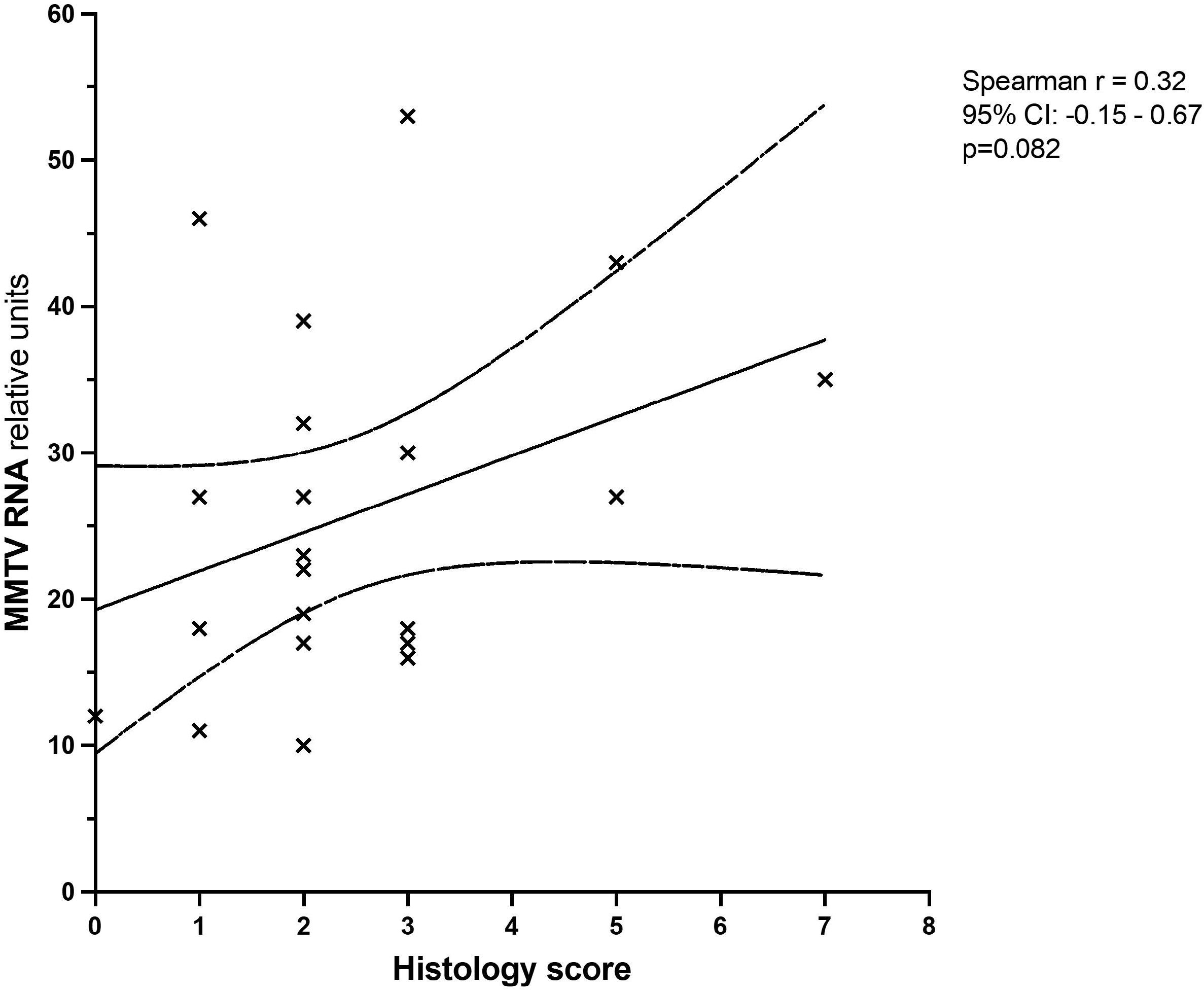
Correlation of colon MMTV RNA levels and histology score [Spearman, p=0.082].

**Supplementary Figure 4:**
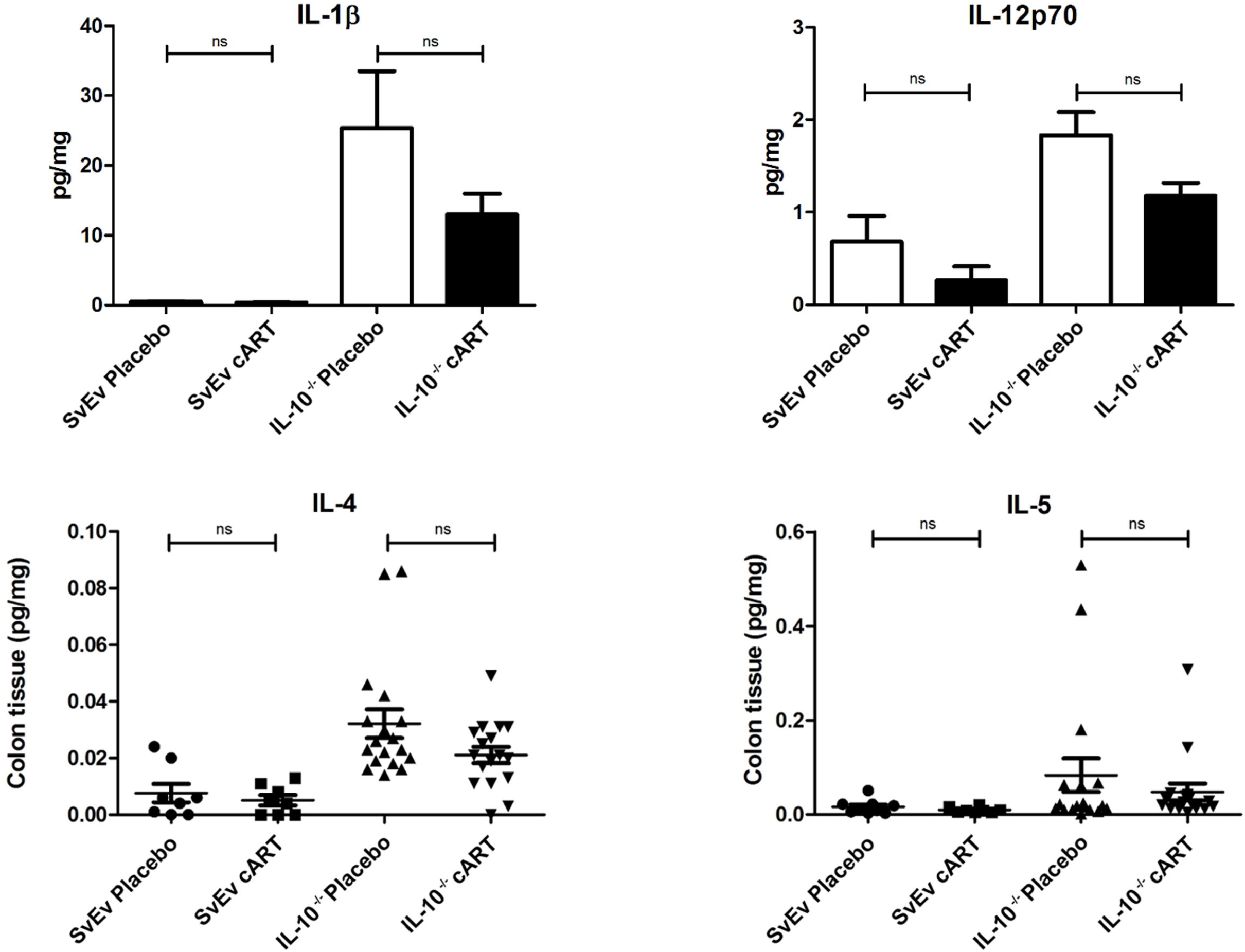
Pro-inflammatory cytokines in colonic extracts showing that levels of IL-1b, IL-12p70, IL-4, and IL-5 were not significantly altered with cART in the IL-10-/- mice [Mean±SEM].

