## Supplementary Table 1 for "Mouse mammary tumor virus is implicated in severity of colitis and dysbiosis in the IL-10^-/-^ mouse model of inflammatory bowel disease"

**Supplemental Table 1**

**Overlapping MMTV Gag peptides**

| **Number** | **Peptide Sequence** |
| --- | --- |
| 1/58 | MGVSGSKGQKLFVSVLQRLL |
| 2/58 | LFVSVLQRLLSERGLHVKES |
| 3/58 | SERGLHVKESSAIEFYQFLI |
| 4/58 | SAIEFYQFLIKVSPWFPEEG |
| 5/58 | KVSPWFPEEGGLNLQDWKRV |
| 6/58 | GLNLQDWKRVGREMKRYAAE |
| 7/58 | GREMKRYAAEHGTDSIPKQA |
| 8/58 | HGTDSIPKQAYPIWLQLREI |
| 9/58 | YPIWLQLREILTEQSDLVLL |
| 10/58 | LTEQSDLVLLSAEAKSVTEE |
| 11/58 | SAEAKSVTEEELEEGLTGLL |
| 12/58 | ELEEGLTGLLSTSSQEKTYG |
| 13/58 | STSSQEKTYGTRGTAYAEID |
| 14/58 | TRGTAYAEIDTEVDKLSEHI |
| 15/58 | TEVDKLSEHIYDEPYEEKEK |
| 16/58 | YDEPYEEKEKADKNEEKDHV |
| 17/58 | ADKNEEKDHVRKVKKVVQRK |
| 18/58 | RKVKKVVQRKEISEGKRKEK |
| 19/58 | EISEGKRKEKDQKAFLATDW |
| 20/58 | DQKAFLATDWNDDDLSPEDW |
| 21/58 | NDDDLSPEDWDDLEEQAAHY |
| 22/58 | DDLEEQAAHYHDDDELILPV |
| 23/58 | HDDDELILPVKRKVVKKKPQ |
| 24/58 | KRKVVKKKPQALRRKPLPPV |
| 25/58 | ALRRKPLPPVGFAGAMAEAR |
| 26/58 | GFAGAMAEAREKGDLTFTFP |
| 27/58 | EKGDLTFTFPVVFMGESDDD |
| 28/58 | VVFMGESDDDDTPVWEPLPL |
| 29/58 | DTPVWEPLPLKTLKELQLAV |
| 30/58 | KTLKELQLAVKTMGPSAPYT |
| 31/58 | KTMGPSAPYTLQVVDMVASQ |
| 32/58 | LQVVDMVASQWLTPSDWHQT |
| 33/58 | WLTPSDWHQTARATLSPGDY |
| 34/58 | ARATLSPGDYVLWRTEYEEK |
| 35/58 | VLWRTEYEEKSKETVQKAAG |
| 36/58 | SKETVQKAAGKRKGKVSLDM |
| 37/58 | KRKGKVSLDMLLGTGQFLSP |
| 38/58 | LLGTGQFLSPSSQIKLSKDV |
| 39/58 | SSQIKLSKDVLKDVTTNAVL |
| 40/58 | LKDVTTNAVLAWRAIPPPGV |
| 41/58 | AWRAIPPPGVKKTVLAGLKQ |
| 42/58 | KKTVLAGLKQGNEESYETFI |
| 43/58 | GNEESYETFISRLEEAVYRM |
| 44/58 | SRLEEAVYRMMPRGEGSDIL |
| 45/58 | MPRGEGSDILIKQLAWENAN |
| 46/58 | IKQLAWENANSLCQDLIRPI |
| 47/58 | SLCQDLIRPIRKTGTIQDYI |
| 48/58 | RKTGTIQDYIRACLDASPAV |
| 49/58 | RACLDASPAVVQGMAYAAAM |
| 50/58 | VQGMAYAAAMRGQKYSTLVK |
| 51/58 | RGQKYSTLVKQTYGGGKGGQ |
| 52/58 | QTYGGGKGGQGSEGPVCFSC |
| 53/58 | GSEGPVCFSCGKTGHIKKDC |
| 54/58 | GKTGHIKKDCKEEKGSKRAP |
| 55/58 | KEEKGSKRAPSGLCPRCKKG |
| 56/58 | SGLCPRCKKGYHWKSECKSK |
| 57/58 | YHWKSECKSKFDKDGNPLPP |
| 58/58 | DKDGNPLPPLETNTENSKNL |
